# Dynamics of *Aedes albopictus* invasion Insights from a spatio-temporal model

**DOI:** 10.1101/2021.09.24.461645

**Authors:** L Roques, T Boivin, J Papaïx, S Soubeyrand, O Bonnefon

## Abstract

France displays a latitudinal range for the expansion of *Aedes albopictus* invasive populations that is not yet completely colonized providing a critical opportunity to address key invasion processes. We propose a spatio-temporal model (DISTIGRI) to describe and predict current and future expansion at both intra- and inter-annual scales of *A. albopictus*. This process-based model integrates mechanistic descriptions of the developmental cycle and the dispersal process of *A. albopictus* within a reaction-diffusion framework, depending on climatic suitability and photoperiod with a high spatio-temporal resolution. Using this model coupled with a climatic database, we propose several maps describing the current intra-annual distribution of *A. albopictus*, including the date of first emergence and the length of the period with significant adult presence. We also compute its future distribution over the next 10 years under several climatic scenarios, which shows a range expansion with a strong dependence on the climatic scenario. The outputs of the model may constitute a valuable asset for designing control and avoidance strategies, and to anticipate the biting nuisance with a high spatio-temporal resolution. These outputs also emphasize the importance of taking both dispersal and life cycle into account to obtain accurate descriptions of out-of-equilibrium processes such as ongoing invasions.

## 1 Introduction

The Asian tiger mosquito *Aedes albopictus* is nowadays considered as one of the most invasive insect species in the world (Global Invasive Species Database, 2021), and is certainly the most invasive mosquito species (Bonizzoni et al., 2013). Native to South-East Asia and to islands of the Western Pacific and Indian Ocean, *A. albopictus* is now present in all continents except Antarctica (Enserink, 2008), where it is a competent vector for many arboviruses (Gratz, 2004) including chikungunya and dengue viruses. *A. albopictus* thus raises major health issues worldwide, and European awareness concerning its introduction and its establishment has increased during the last decade with subsequent transmission and outbreaks of the chikungunya virus in Southern Europe (Lindh et al., 2019). Besides its pathogen transmission ability, *A. albopictus* is an aggressive daytime biter and induces a significant nuisance, with important social and economic impacts (Darbro et al., 2017).

The global spread of *A. albopictus* mainly results from increased globalization involving intercontinental air travel and global shipping transport (Paupy et al., 2009). Its use of container habitats for breeding (e.g. used tyres and water containers in domestic settings) has driven its international spread and establishment close to human habitations (Ali and Nayar, 1997; Benedict et al., 2007). In general, the incidence and geographic range of *Aedes* species is now expected to expand dramatically in response to urbanization and socio-economic factors, but also environmental changes such as warmer global temperatures (Patz et al., 1998; Messina et al., 2015; Kraemer et al., 2019). In Europe, it was first detected in Albania in 1979 (Adhami and Reiter, 1998), and it spread to all Mediterranean countries within about two decades (Caminade et al., 2012) through combined long-range dispersal events (e.g. in tyres and potted plants) and local diffusive movements facilitated along roads (Roques and Bonnefon, 2016). In France, the first record of *A. albopictus* was in 1999 (Schaffner and Karch, 2000) in the northern part of the country, whose cold climate probably prevented the establishment of an invasive population. The first established population of *A. albopictus* in France was detected in 2004 in south-eastern France, with a likely Italian origin (Scholte and Schaffner, 2007), and invasive populations are now gradually expanding northwards as elsewhere in Europe (e.g. Caminade et al., 2012; Medlock et al., 2012; Roques and Bonnefon, 2016). The overall relatively low rate of spread of *A. albopictus* at northern latitudes in Europe likely supports the importance of climate conditions on the establishment and spread of the species.

The effects of climate and photoperiod on the life history of *A. albopictus* are now well documented, through laboratory experiments (Delatte et al., 2009), *in natura* observations (Kobayashi et al., 2002; Toma et al., 2003) and climatological analyses. The ecological plasticity of *A. albopictus* may explain successful expansions towards northern cooler climates. In particular, *A. albopictus* is able to induce photoperiodic egg diapause, allowing overwintering survival in temperate regions (Hawley, 1988) and population establishment at higher latitudes than other exotic mosquito species that do not produce diapausing eggs. As European winter temperatures still remain an ecological limiting factor for survival (Medlock et al., 2006; Caminade et al., 2012) and as spring and summer climate conditions play a key role on larval development (Komagata et al., 2017) and adult reproduction (Delatte et al., 2009), it is acknowledged that climate change is a key driver of expansion towards northern latitudes (Caminade et al., 2012). In this context, one might expect important variations in *A. albopictus* dynamics along latitudinal gradients, and the sanitary, social and economic impacts of *A. albopictus* on human activity raise major issues in forecasting how intra-annual mosquito dynamics at any given location might change due to both expansion processes and environmental change.

Models are paramount tools for predicting the response of *A. albopictus* populations to environmental change. In their review, Fischer et al. (2014) classified the existing models as correlative niche models and mechanistic models. In correlative models, the niche of *A. albopictus* is determined by statistical correlation between observations (e.g., presence and/or absence) and climatic variables. For instance, the European Centre for Disease Prevention and Control (2009) used random forest models to assess the climatic suitability of Europe for *A. albopictus*. Mechanistic models fit into two classes depending on whether the models are process-based or not. Mechanistic approaches that are not process-based rely on the construction of overlay functions for climatic constraints in a geographic information system. These approaches do not describe the life cycle of the mosquito and its dispersal but rely on suitability criteria. Using this type of approach, Medlock et al. (2006) evaluated the ability of *A. albopictus* to establish in the UK and its seasonal activity. Caminade et al. (2012) used comparable approaches to assess the suitability of Europe for *A. albopictus* using both recent and future climate conditions. Process-based approaches incorporate the life cycle of the mosquito and its interactions with the environmental constraints such as climate, photoperiod and landscape features. These approaches often rely on compartmental models, describing the different stages of the life cycle. Existing approaches either do not include dispersal (ODE, ordinary differential equation models) or are rather theoretical reaction-diffusion models (Tran et al., 2013; Strugarek et al., 2019; Haramboure et al., 2020; Tran et al., 2020; Dufourd and Dumont, 2012, 2013; Roques and Bonnefon, 2016). At the European scale, Fischer et al. (2014) emphasized that none of the existing modelling studies on expected future abundance of *A. albopictus* explicitly incorporated processes such as the introduction and dispersal of the mosquitoes. To the best of our knowledge, this is still the case nowadays.

In this study, we took the opportunity that France displays a critical latitudinal range for *A. albopictus* expansion that is not yet completely colonized to develop a single modelling approach involving the intertwined effects of dispersal and climate on *A. albopictus* distribution. For this purpose, We develop here a reaction-diffusion compartmental model that takes into account both temperatures and photoperiod data at a high spatial and temporal resolution, and that explicitly describes *A. albopictus* adult dispersal over the French territory, with faster dispersal along the main roads. The main objectives of our DISTIGRI model (DISpersal of TIGeR mosquIto) are: (1) to estimate the distribution of key variables such as the date of the first adult emergence each year and the duration of the period with significant adult occurrence, depending on the spatial position in the country; (2) to provide a tool for a real-time monitoring of the intra-annual dynamics of *A. albopictus* depending on the spatial position and on the previous and current climate conditions; (3) to forecast the expansion of *A. albopictus* over the next 10 years, with several climatic scenarios; (4) to provide an updated assessment of the climatic conditions that drive the winter survival and adult emergence. We expect that such modelling outcomes can help improve the targeted monitoring of *A. albopictus* populations and guide the implementation of controlling programs to prevent nuisance and further expansion.

**Figure 1:**
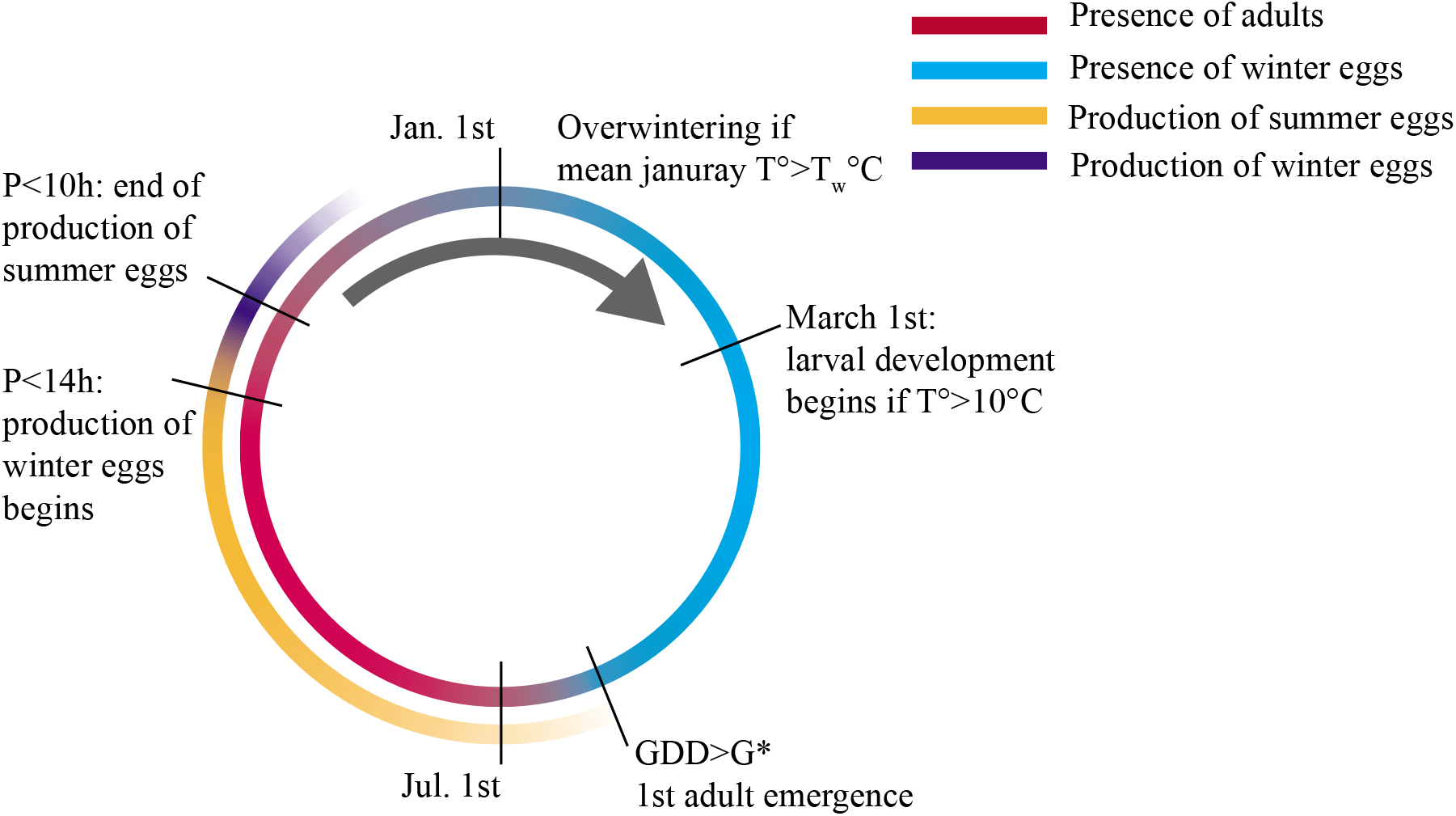
Schematic life cycle of *A. albopictus* over one year in France. *P* corresponds to the photoperiod duration (in hours), and GDD is the growing degree days (in °C days) since March 1^st^, see equation (1).

## 2 Material and methods

### 2.1 Life cycle and construction of the DISTIGRI model

We describe here the aspects of the life cycle of *A. albopictus* that are integrated in the DISTIGRI model (see Fig. 1 for a synthetic view) together with the associated modelling tools.

#### First adult emergence

The date of first adult emergence depends on late winter and spring temperatures. Hawley (1988) observed that larval development ceases at temperatures below 11°C. Based on this observation, several studies assumed that hatching starts when spring temperatures are above a threshold comprised between 10 and 11°C, see Model 3 in Caminade et al. (2012), or the report of the European Centre for Disease Prevention and Control (2009). In other approaches, instead of a threshold value at current time, mean spring temperatures are used (Medlock et al., 2006). In order to match with the biological processes underlying the hatching date, we assume here that the emergence of the first adults depends on the cumulated difference between daily mean temperatures at their position, and a threshold value for development. This assumption is based on the recent work of Komagata et al. (2017) who studied the association between the first biting day and spring temperature in several locations of Japan. They computed a growing degree days (GDD) value, based on the cumulated difference between daily mean temperatures and a threshold value for development of 10°C:

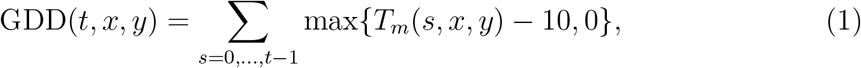

with *t* the number of days since March 1^st^, (*x, y*) the spatial position, and *T_m_* the daily mean temperature. Komagata et al. (2017) obtained a mean GDD of 269°C days, with a 95% confidence interval (246, 294).

In our approach, adult emergence begins at some date *t*_0_(*x, y*) when the GDD reaches a threshold *G*\*(*G*\* =246, 269 or 294°C days, see below). It occurs at a rate *γ*_0_(*t, x, y*), which is equal to 0 before *t*_0_. After the date *t*_0_, we assume a constant rate of adult emergence: *γ*_0_(*t, x, y*) = *h*_0_, for *t* > *t*_0_(*x, y*). In Toma et al. (2003), a mean hatching rate of about 20% per period of 2 days was recorded at a mean temperature of 12.3°C. Consistently with this observation, we assume that *h*_0_ is such that *e*^−2*h*_0_^ = 0.8, i.e., *h*_0_ ≈ 0.1. The density *w*_0_(*t, x, y*) of diapausing winter eggs at the beginning of the year then obeys the following differential equation:

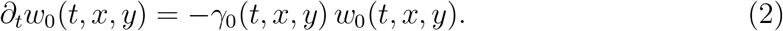

We neglect the effect of the photoperiod on the occurrence of the first adult emergence, as several studies indicate that the photoperiodically sensitive stages are the adults and pupae (Wang, 1966; Imai and Maeda, 1976; Mori et al., 1981). The photoperiod should therefore not affect much the development of diapausing eggs.

#### Adult stage and production of summer eggs

After emergence, the adults disperse and reproduce, leading to new ‘summer’ eggs which in turn produce new adults. The ovo-larval stage and the mating and feeding of the adults are not modelled explicitly but taken into account through a growth term *R*(*t, x, y, a*), with *a* = *a*(*t, x, y*) the adult population density. The growth term depends on the current temperature *T*(*t, x, y*) and photoperiod *P*(*t, x, y*):

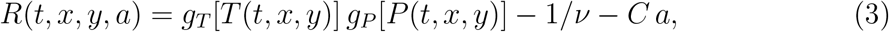

with *ν* the life expectancy of the adults in the absence of competition (30 days, see Delatte et al., 2009), *g_T_, g_P_* two functions which measure respectively the effect of the temperature and of the photoperiod on the growth rate. The calibration of the function *g_T_* is based on the laboratory experiments described in Delatte et al. (2009), see Supplementary Information. Regarding the effect of the photoperiod, we assume that, during the first part of the year (before June 21^st^), *g_P_*(*P*) = 1 (no effect of the photoperiod). Then for the second part of the year, (after June 21^st^, when the photoperiod starts to decline), based on the observations of Toma et al. (2003), we assume that the production of summer eggs starts decreasing once a critical photoperiod of 14h is reached, and then decreases linearly until the photoperiod reaches 10h, where the production of summer eggs stops (*g_P_* = 0). The term ‘*Ca*’ is a density-dependent mortality term; the parameter *C* is such that, in optimal conditions (*g_T_* = max*g_T_*, *g_P_* = 1), the positive steady state (a such that *R*(*t, x, y, a*) = 0) is equal to the carrying capacity *K*. Without loss of generality, we take *K* = 1; this means that the number of adults *a* is expressed in units of the carrying capacity.

The adult stage is then modelled with a reaction-diffusion equation. To take into account the effect of roads on the dispersal of the adults we adopt the framework developed in Roques and Bonnefon (2016). This framework is based on a coupled system of 2D (dynamics in patches) and 1D (dynamics on roads) equations. For the sake of simplicity, we only describe here the population dynamics in the 2D patches (the full model that we used in our simulations is available as Supplementary Information):

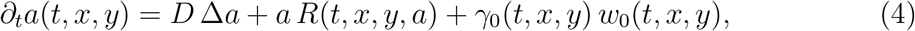

with *D* the diffusion coefficient, which measures the mobility of the adults, assuming a random walk movement which is described by the Laplace differential operator Δ, (see e.g. Turchin, 1998).

#### Production of winter eggs

Once the photoperiod of 14h has been reached, the adults start to produce diapausing winter eggs which will only hatch during the next spring. In the DISTIGRI model, the winter eggs replace the summer eggs, i.e., the number of winter eggs that are produced correspond to the number of summer eggs that are not produced, due to diminished photoperiod. This leads to a rate of production of winter eggs:

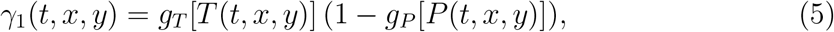

and the density *w*_1_ of winter eggs follows the differential equation:

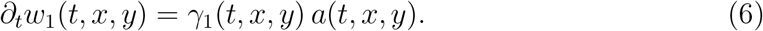

#### Overwintering and inter-annual stage

At present, it is usually accepted that winter eggs survival in Europe depends on the mean January temperature, with a lethal threshold of 0°C (Burgess, 1995; Medlock et al., 2006; European Centre for Disease Prevention and Control, 2009). As Caminade et al. (2012) (Model 1) we consider here three different thresholds in this study *t_w_* = 0°C, 1°C or 2°C. The density of diapausing winter eggs at the beginning of the year *n* (March 1^st^), 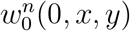 depends on the density of diapausing winter eggs at the end of the year *n* – 1, 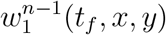, with *t_f_* corresponding to December 31^st^ of year *n* – 1. This dependence is modelled as follows:

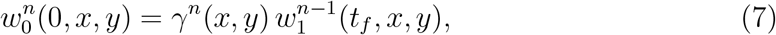

with *γ^n^*(*x, y*) a parameter describing the winter survival rate. This parameter depends on the mean January temperature of year *n*, 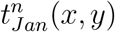, as follows: *γ^n^*(*x, y*) = 0 if 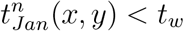 (no overwintering) and *γ^n^*(*x, y*) = 1 (no mortality) otherwise.

#### Overwintering and adult emergence: 3 levels of risk

We consider 3 levels of risk in terms of human exposure to mosquito biting, depending on the GDD threshold *G*\* that determines adult emergence, and of the overwintering threshold *t_w_*: (i) low risk: *G*\* = 294° days, *t_w_* = 2°C; (ii) intermediate risk: *G*\* = 269° days, *t_w_* = 1°C; (iii) high risk: *G*\* = 246° days, *t_w_* = 0°C.

#### Numerical simulation

The simulations were carried out using a finite-element method in space and an implicit scheme in time. The non-linearity was dealt with using a Newton-Raphson algorithm. The simulations were performed using the Freefem++ finite-element framework (Hecht, 2012).

### 2.2 Data

#### Study site

The study site is mainland France. As mentioned above, following the framework developed in Roques and Bonnefon (2016), in addition to standard 2D diffusion, we take into account the effect of the main roads on the propagation of the mosquito. For simplicity, we consider only the main highways, with an average daily traffic of 15,000 vehicles or more (source: French Centre For Studies and Expertise on Risks, Environment, Mobility, and Urban and Country planning, CEREMA, 2011).

#### *A. albopictus* distribution: inter-annual data

The first established population was detected in 2004 in the Alpes-Maritimes county (the South-Easternmost French county: mainland France is segmented in 94 administrative units called ‘départements’), probably as a result of expansion of the Italian insect population (Scholte and Schaffner, 2007). Introductions may have also occurred from Northern Spain, where it has been reported since 2004 (Aranda et al., 2006). In this study, we use the 2008-2019 data, which were available from the European Centre for Disease Prevention and Control, see Supplementary Fig. S2. At the scale of a French county, three levels of infestation have been defined by the ECDC: ‘absent’, field surveys were conducted and no introduction or no established population of the species have been reported; ‘introduced’: the species has been observed (but without confirmed establishment) in the administrative unit; ‘established’: evidence of reproduction and overwintering of the species has been observed in at least one municipality within the administrative unit.

In each county of mainland France, we define an observation variable 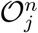, which depends on the status of *A. albopictus* in the *j*^th^ county during year *n*:

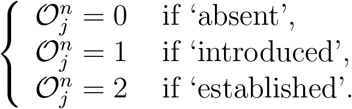

This corresponds to a total of 12 × 94 = 1128 observations.

#### Climatic data

The temperature data at the hourly time step on a 8 × 8km^2^ grid covering France were provided by the SAFRAN database over the period ranging from August 1, 2008 to July 31, 2019. SAFRAN is based on an interpolation method of climate variables measured at ground level. Both observations from meteorological stations of the French national weather service and surface analyzes from numerical weather prediction systems are used. For more details about SAFRAN, see Durand et al. (2009).

#### Simulated climate scenarios

We constructed a simulated climatic dataset as follows. Each year *n* in 2020-2030 the climate data are selected at random from the 2008 – 2019 data (full year, with replacement). With this algorithm, we built 100 independent climate series to take into account the climate variability and the uncertainty about future climate trends. We discuss the simulation of climate scenarios in the Discussion section.

#### Initial state

We construct the initial condition as follows. First, at September 1^st^ of year *n*_0_ – 1, we assume the following adult densities: *a* = 0.5 in the ‘established’ regions, *a* = 0.25 in the ‘introduced’ regions, and *a* = 0 in regions where *A. albopictus* is considered as absent (see the observation model in Section 2.3 for a justification of these values). Then, we run the dynamical model until March 1^st^ of year *n*_0_. This leads to a density 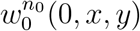 that we use as the initial condition of the model.

### 2.3 Model validation

#### Observation model

In order to compare the continuous adult densities *a*(*t, x, y*) given by the DISTIGRI model with the discrete observations data 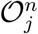 of Section 2.2, we develop a probabilistic approach, which describes the observation process conditional on the adult mosquito density. This approach is inspired from the framework of Roques and Bonnefon (2016) which relies on the mechanistic-statistical theory (e.g., Ovaskainen et al., 2008; Soubeyrand et al., 2009; Soubeyrand and Roques, 2014).

Namely, for each county, we define 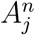 the maximum (over space and time) adult density in the county *j* during year *n*. Then, we assume that the observation in the corresponding county is drawn from a binomial distribution, conditional on 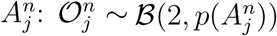. The function *p* describes the relationship between 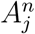 and the probability of mosquito detection. For the sake of simplicity, and because the maximum adult densities that we obtained in our simulations are ≈ 0.5 (with *K* =1, which would be reached under optimal conditions), a natural choice for *p* is

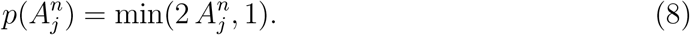

*Note*. To check that this modelling approach gives reasonable results, we detail below the distribution of the observations, for some particular values of the maximum adult density *A*. We denote by 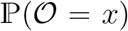 the probability of the event 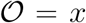, for *x* = 0,1 or 2. If *A* ≥ 0.5 then 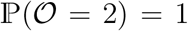, which means that the detection of established populations in the only possible outcome; if *A* = 0.25 then 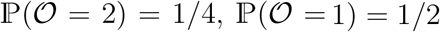 and 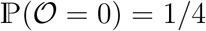, i.e., the detection of introduced populations is the most probable outcome; if *A* = 0, then 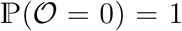, and the only possible observation is ‘absent’.

#### Assessment of the mean predictive power at the inter-annual scale

To assess the goodness of fit and predictive power of the dynamical model, we use the observation data of the year *n*_0_ – 1 to build the initial condition *w*^*n*_0_^ and we check if the model gives accurate predictions at later years. We use the multi-category half-Brier score *BS* (Gneiting et al., 2007), a mean squared error measure of probability forecasts which is well-adapted to discrete observation data. The score *BS* = 0 corresponds to a perfect forecast and *BS* =1 corresponds to the worst score. With data including three categories (0=absent, 1=introduced and 2=established), we define the *k*-year predictive power starting from the year *n*_0_ by:

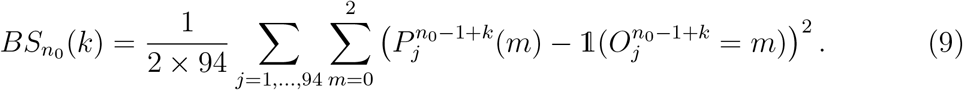

Here, 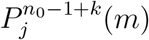 is the ‘forecast probability’, i.e., the probability that event *m* occurs during the year *n*_0_ – 1 + *k* in the county *j* with the above observation model and based on the output of the dynamical model. Namely, we define as above 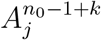 the maximum (over space and time) adult density in the county *j* during year *n* given by the dynamical model. With 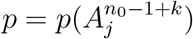 (see eq. (8)), we deduce from the binomial assumption on the observations that 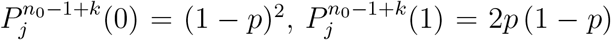, and 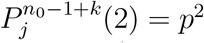.

For *k* = 1 (1-year predictive power) we compute *BS*_*n*_0__(1) for each value of *n*_0_ between 2009 and 2019. We denote by 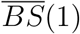 the mean value of *BS*_*n*_0__ (1) over these 11 years. Similarly, for *k* larger than one 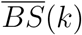 is the mean *k*-year predictive power, averaged over the 12 – *k* admissible years *n*_0_ = 2009,…, 2019 – *k* + 1.

We compare these values with the half-Brier score obtained with a reference model that we denote by 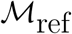. In this reference model, we assume that the population remains in a constant state, corresponding to the observation of year *n*_0_. More precisely, we assume that the adult density *a* is 0.5 in the ‘established’ regions of year *n*_0_, 0.25 in the ‘introduced’ regions of year *n*_0_, and 0 in regions where *A. albopictus* is considered as absent during year *n*_0_, and that these densities do not change at further years *n*_0_ + *k*.

## 3 Results

### Model validation

We computed the mean *k*—year predictive power for the three scenarios, with a maximal horizon *k* = 11. The results are depicted in Fig. 2. We note a remarkably stable value of the mean Brier score as the horizon *k* is increased. In particular, it outcompetes the reference model after 3 years, and the difference of accuracy between the two models increases as we increase the horizon of prediction *k*. For instance, after 5 years the mean Brier score obtained with the reference model is 58% higher, compared to the DISTIGRI model with the ‘high risk’ parameter values. Due to its definition, the Brier score intrinsically gives an advantage to the reference model. In particular, in the regions where *A. albopictus* is considered as being absent, the adult density is set to 0 in 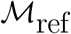, whereas although possibly small, the density is never exactly equal to 0 with the DISTIGRI model. A similar phenomenon can occur in the regions where *A. albopictus* is considered as being established. Thus, in the counties that do not change (i.e., which keep the same color in the data of Supplementary Fig. S2), the Brier score is always larger with the reference model. This explains the higher 1- and 2-year predictive powers obtained with the reference model, technically advantaged when the distribution of *A. albopictus* remains stable.

Regarding the effect of the parameters, we note that the mean Brier score is slightly improved as we move away from the ‘low risk’ parameters. Thus, the values *G*\* = 246° days and *t_w_* = 0°C corresponding to the high risk parameter values seem more appropriate for the dynamics of *A. albopictus* in France over the previous decade. We use these values in the subsequent computations.

**Figure 2:**
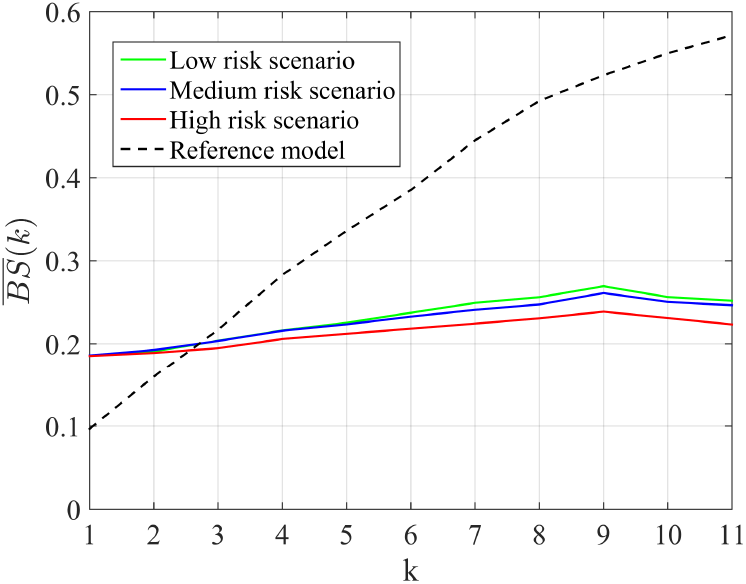
Assessment of the predictive power: mean Brier score at *k* years.

### Intra-annual dynamics

We computed the intra-annual dynamics in three cities in France, Marseille, Toulouse and Paris, with contrasted climates (Mediterranean, temperate humid and oceanic, respectively) and different dates of first colonization (2009, 2012 and 2015, respectively, based on the data in Supplementary Fig. S2). The results are presented in Fig. 3, and the positions of the cities are visible on Fig. 4. In all cases, the maximum adult density is larger in 2018, a hot year compared to 2017 (mean annual temperatures: 13.4°C in 2017 and 13.9°C in 2018, source: Météo France). This also leads to a larger number of overwintering eggs at the end of 2018. As expected, the duration of the period where adults are present with a significant density (adult population density *a* > 0.1) varies with the city and with the year: in 2017 this duration is of 139 days in Marseille, 130 in Toulouse and 0 in Paris. In 2018: 150, 136 and 72 days respectively. The variability between the two years is higher in Paris. This reflects the recent introduction in this region and the undergoing expansion process, compared to Marseille and Toulouse.

**Figure 3:**
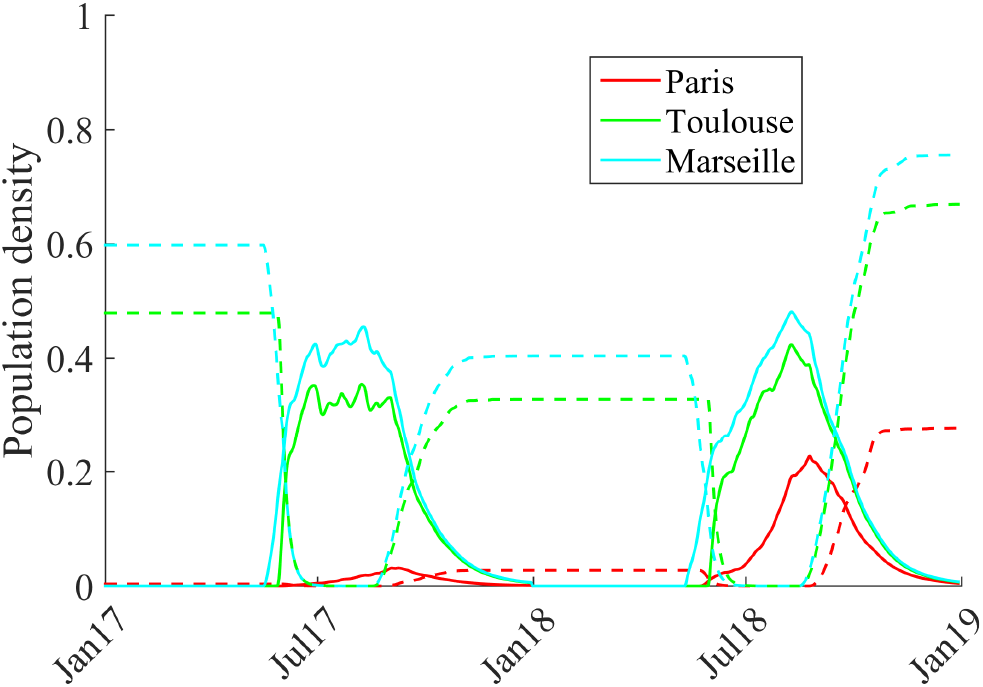
Intra-annual dynamics of the egg and adult density in three cities over two years. The plain lines correspond to the adult density *a* and the dotted lines to the sum of the summer and winter egg densities, *w*_0_ + *w*_1_.

We depict in Fig. 4 the theoretical dates of first emergence *t*_0_ (*x, y*) and the duration of the period with significant adult presence (defined here as the length of the period where *a* > 0.1) in 2017, 2018 and 2019. We observe important changes in *t*_0_ between these three years, with an earlier emergence in 2018 compared to 2017 and 2019. This variability between years and over the territory is the consequence of variability in the climate conditions, as the date of first emergence only depends on the GDD (growing degree days) value at the beginning of each year. On the other hand, the duration of the period with significant adult presence not only depends on the climate conditions, but also on the previous presence of the mosquito and on its dispersal. We indeed observe in Fig. 4 that it keeps increasing from 2017 to 2019 despite much later emergence in 2019. We also observe an earlier invasion of the Paris regions compared to surrounding regions with similar *t*_0_ values. This early invasion is probably caused by the faster dispersal on the main roads, that we take into account in the model. For the same reason, we observe in 2019 a significant presence of the adults along the main roads around the Paris Basin.

We compare in Fig. 5 the portion of the territory where overwintering was possible vs the regions with a significant presence of adult mosquitoes (regions where the mean adult density in August exceeds 0.1) in 2017, 2018 and 2019. We observe important fluctuations of the overwintering area, which occupied 64%, 97% and 86% of the territory from 2017 to 2019. In spite of these fluctuations, the area with a significant presence of adult mosquitoes keeps increasing, and occupies about 38% of the territory in 2019, which also means that the potential climatic niche is not yet entirely colonized. Interestingly, adults can be present by dispersal in regions slightly outside the overwintering area (this is visible in 2017 and 2019, but not in 2018, where the overwintering area occupies almost all of the territory).

**Figure 4:**
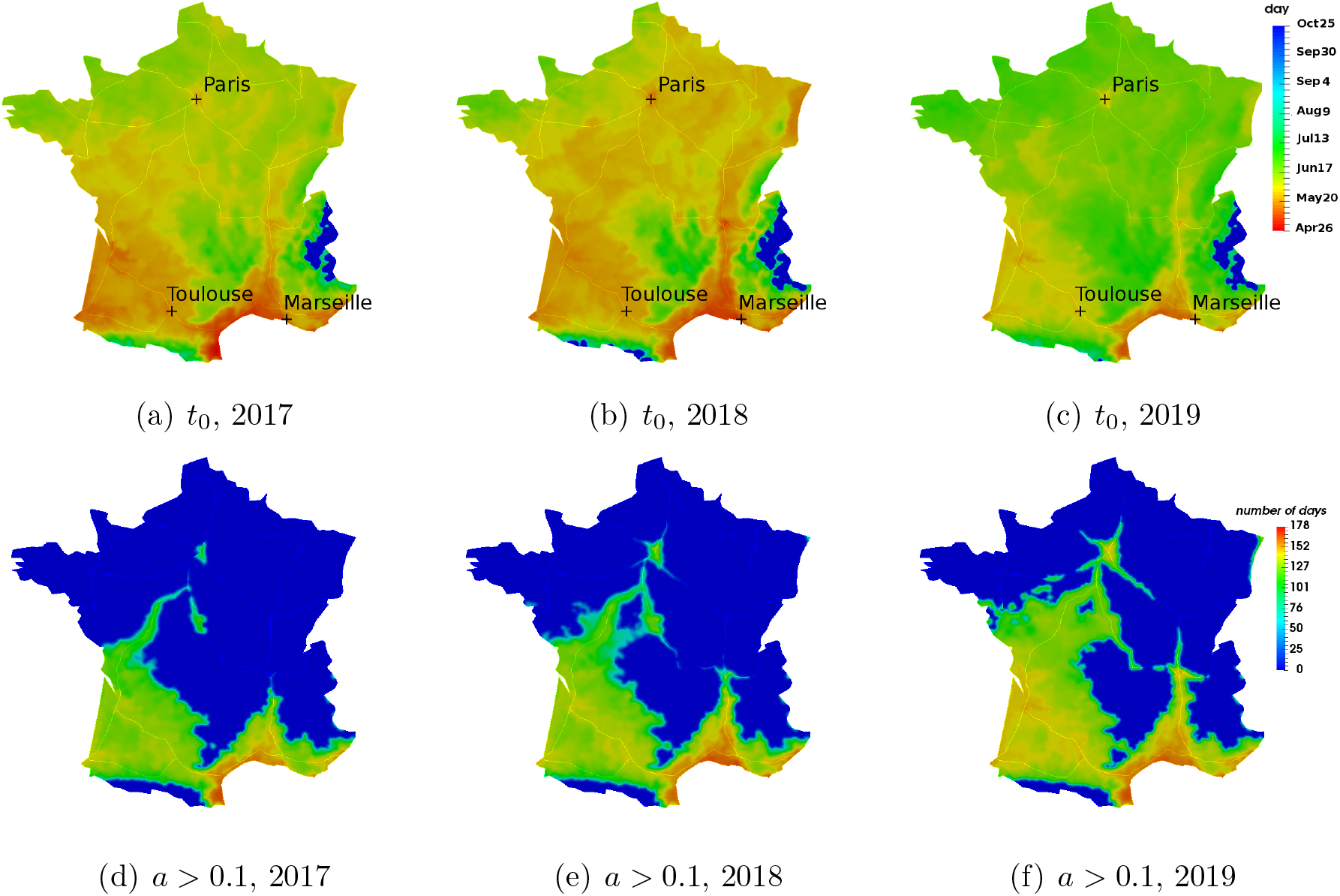
Theoretical date of first adult emergence (a,b,c) and duration of the period with significant adult presence (d,e,f). At each position (*x, y*), the date *t*_0_(*x, y*) depicted here corresponds to the moment when the growing degree days (GDD, see formula (1)) reach the threshold *G*\* = 246 days (high risk parameter values). The duration of the period with significant adult presence is defined here as the length of the period where *a* > 0.1.

### Predictions

We depict in Fig. 6 the predicted mean adult population density in August (the adult population is expected to approach its highest values in August), over the period 2021-2030. We show the 0.05-quantile (corresponding to a cold scenario), the median value (temperate scenario) and the 0.95-quantile (warm scenario), obtained from 100 independent simulated climate series.

We note an important effect of the latitude and of the climate scenario. For each fixed quantile, we observe a spatial expansion of the species range. From a quantitative viewpoint, the cold scenario (0.05-quantile) leads to a significant presence of adult mosquitoes (regions where the mean adult density in August is above 0.1) over 36% of the territory in 2021, 40% in 2024, 44% in 2027 and 41% in 2030. In the temperate scenario, we obtain 47% of the territory in 2021, 51% in 2024, 55% in 2027 and 56% in 2030. In the warm scenario (0.95-quantile), 60% of the territory in 2021, 67% in 2024, 73% in 2027 and 74% in 2030. The increase in the proportion of the occupied territory is mostly due to an expansion in Northern France (the unoccupied areas in the South mostly corresponding to cold mountain areas).

**Figure 5:**
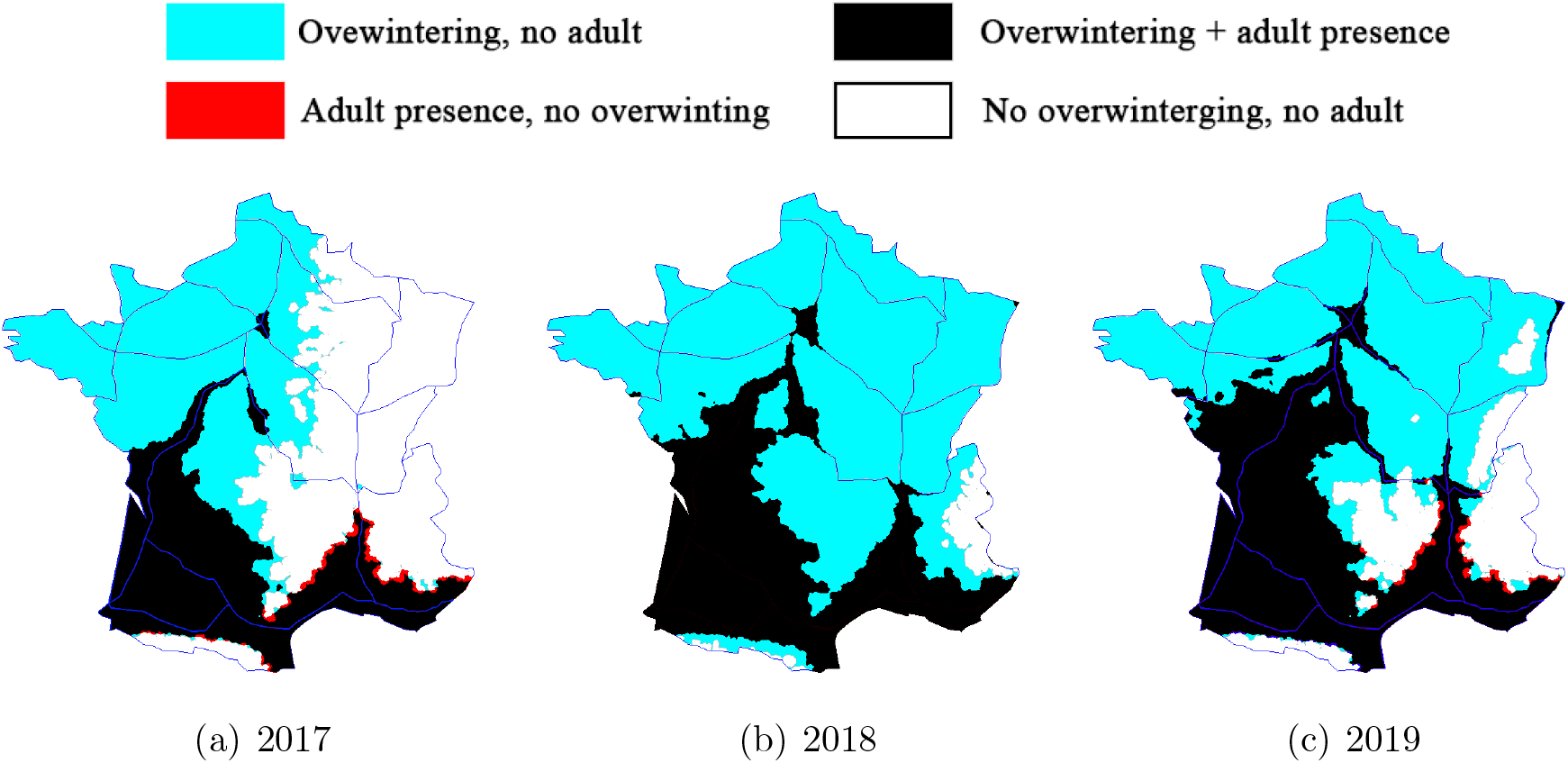
Regions where overwintering is possible vs actual presence of adult mosquitoes. In the blue regions and black regions, overwintering is possible (mean January temperature above 0°C). In the black regions and red regions, the mean adult density in August is higher than 0.1.

In the cold and temperate scenarios, the predicted expansion is likely not driven by increasing temperatures. It is rather a consequence of an increased occupancy of the overwintering region, due to adult dispersal and reproduction. Still, the important difference between the temperate and warm scenarios indicate that increasing temperatures would have a significant effect on future distribution. In particular, in the warm scenario, overwintering is presumably not a limiting factor to range expansion. We note that, in the cold scenario, the expansion slows down and stops from 2027, probably due to a saturation of the climate niche, while it continues in the other scenarios though at a lower rate.

The predicted length of the period with significant adult occurrence is available as Supplementary Material (Supplementary Fig. S4). It shows comparable trends as the adult population density in Fig. 6, but is of independent interest as a proxy of the biting nuisance.

**Figure 6:**
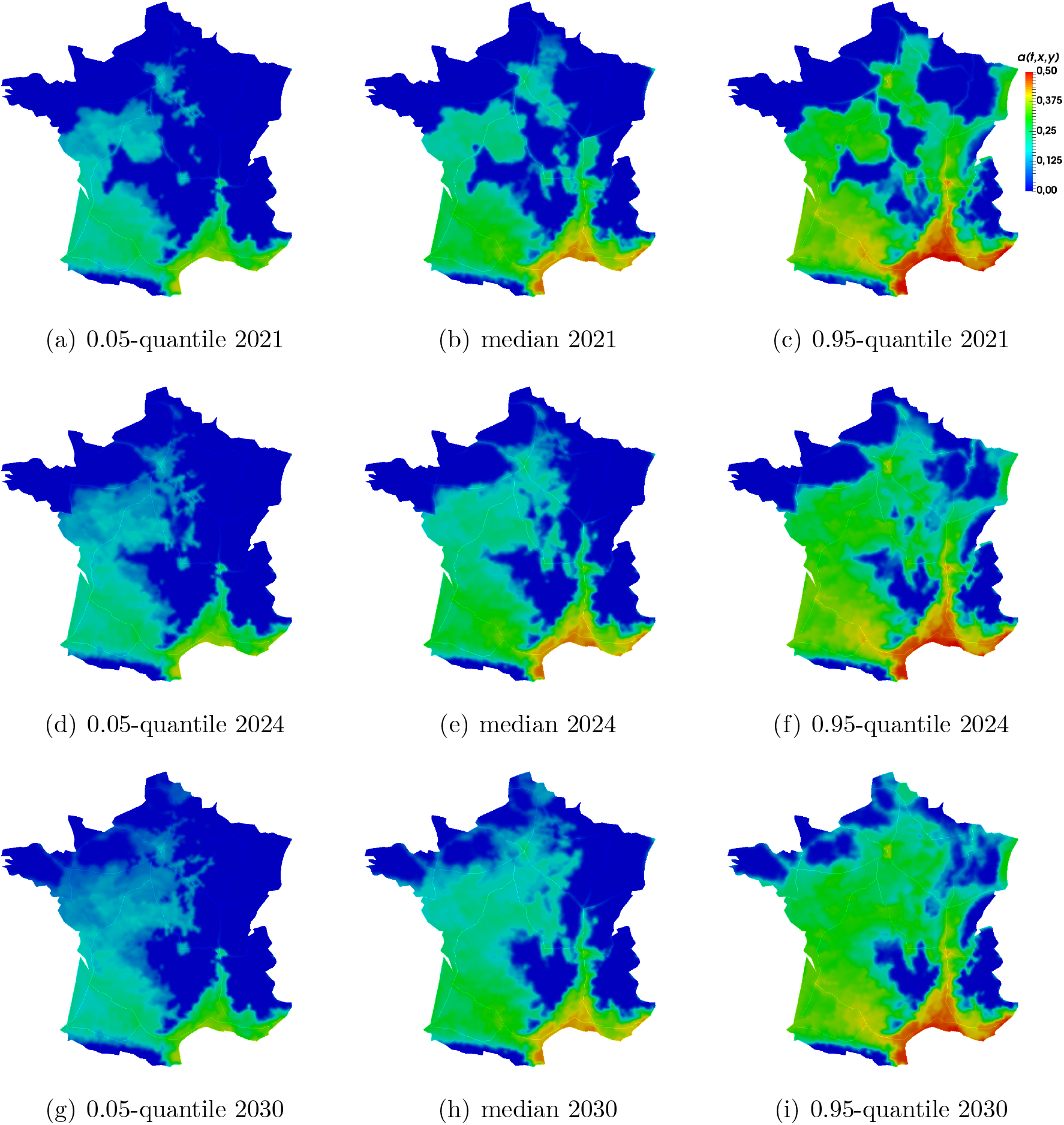
Predicted adult density in August over the period 2021-2030. At each position (*x, y*) we show the 0.05-quantile, the median value and the 0.95-quantile of the average adult population density in August (left, central and right columns, respectively). The distribution and the quantiles are computed from 100 replicate simulations with random climatic data (selected from the 2008 — 2019 data, full year). The predictions for 2027 are available as Supplementary Fig. S3.

## 4 Discussion

We computed the current distribution of *A. albopictus* and its future distribution over the next few years, under various climate scenarios. The DISTIGRI model not only takes the climatic suitability into account but also integrates a mechanistic description of the developmental cycle and of the dispersal process. At a global scale, it predicts an expansion from about 38% of the territory in 2019 to 56% in 2030, with important variations depending on future climate trends (90%-IC: 41%-74%). At a local scale, it gives estimations of the date of first emergence and of the length of the period with significant adult presence. The corresponding outputs of the model, including the maps that are provided in this work, may constitute a valuable asset for designing control and avoidance strategies, and to anticipate the biting nuisance with a high spatio-temporal resolution.

More generally, our results show the importance of taking the dispersal and life cycle into account to obtain accurate descriptions of ongoing invasion processes. With correlative niche models such as species distribution models, or mechanistic approaches that do not consider these processes (e.g. the models discussed in Fischer et al., 2014), the population would remain stable under constant climate conditions. Climate-based predictive models rely on straightforward assumptions when the link between species distribution and climate is well known, but their examination require caution in most cases as non-climatic drivers of species distributions may be important and make prediction challenging (Guo, 2003; Sexton et al., 2009). In a few exceptions, correlative models used non-climatic predictors such as land cover (Thuiller et al., 2004; Ay et al., 2017), biotic interactions (Heikkinen et al., 2007; Palacio and Girini, 2018) and dispersal (Boulangeat et al., 2012). Thus, it remains crucial for invasion models to decipher the relative influence of climate and other ecological factors on species expansion. Here, the model predicts an expansion towards North even in the absence of climatic changes (median ‘temperate’ climatic scenario). This shows that dispersal is still a key limiting factor for invasions as all potentially suitable areas may not be colonizable, and in our case the entire climatic niche is not yet colonized. Conversely, at the intra-annual scale, adults can be present by dispersal in regions outside of the climate niche. This cannot be directly predicted by correlative models, but likely reflects the real situation: theory on the relationship between niche and distribution admits that a species may be absent in some suitable regions and present in unsuitable regions (Pulliam, 2000). This is especially the case during invasions, as invasive species are not in equilibrium with their environment (Gallien et al., 2012), leading to misinterpretation of the realized climatic niche. Regarding that aspect, our consideration of dispersal and of the intra-annual life cycle can greatly improve projections. Correlative niche models thus give a first approximation of the potential presence of the species at a lower computational cost, where process-based approaches that explicitly describe the dispersal and reproduction of the individuals (and possibly interactions with other species) are able to predict the realized occurrence of the species.

On the other hand, climate remains an important driver of the expansion of *A. albopictus*, as a shift toward higher temperatures would lead to an important change in the proportion of the territory with a significant presence of adult mosquitoes. Although current climate projections predict warmer conditions in Europe (Meehl et al., 2007), the observations in 2017 (Fig. 5) show that one may also expect strong fluctuations of the overwintering area, that could lead to transient range retractions even with a global tendency to increasing temperatures at the annual scale.

We observed that the high risk parameter values gives more accurate results. In the literature, the lethal mean January temperature is generally between 0 and 2°C. These values are based on observations in Japan (Medlock et al., 2006). The threshold growing degree days (GDD) value also corresponds to observations in Japan (Komagata et al., 2017). Generally speaking, invaded climatic niches may however differ from native climatic ones so that the species realized niche is not conserved over space (Early and Sax, 2014; Barbet-Massin et al., 2018). Regarding *A. albopictus*, Medley (2010) showed a niche shift from its native state during invasion of both America and Europe. This suggests that niche conservatism doesn’t apply to the spread of invasive *A. albopictus* populations, and that parameter values inferred from the native regions should be updated. Our study leads to such updated values, in a European country. We did not try however to test the model with even lower thresholds (corresponding to higher risk). There is a possibility that the overwintering threshold is strictly below 0°C, as some observations in China and South Korea indicate (Nawrocki et al., 1987). With the same framework, we could use the Brier Score to obtain a more accurate estimation of the parameters, although this optimization procedure would be computationally costly.

In this study, we assessed the potential expansion of *A. albopictus* at a relatively short temporal horizon (10 years). Hence, we applied a simple resampling technique to generate future annual climates from observed annual climates (stochastic weather generators could be used as well, to generate completely new annual climates statistically distributed like observed climates; Allard et al., 2015). For longer temporal horizons, we should account for global-warming in our forecast by using climatic projections grounded on different representative pathways of CO2 concentration (e.g., obtained from the DRIAS platform; http://www.drias-climat.fr/). Since immature stages are aquatic, one could also expect that the species relies on regular rainfalls for breeding. However, in modern urbanised countries such as France, *A. albopictus* has the ability to breed in places that are independent of rainfall, such as urban gardens, barrels, rainwater gulley catch basins and drinking troughs (Gatt et al., 2009). As underlined by Waldock et al. (2013), the effect of rainfall is limited in such permanent water sources, and rainfall mostly affects the creation of additional breeding sites, with overall a variable effect of rainfall on observed population densities. This makes the effect of precipitation one of the most difficult to model. The usual criterion for survival is that the annual rainfall is above 500mm (Medlock et al., 2006), a value which is reached in all regions of mainland France according to the French official meteorological and climatological service (Météo France). The presence of *A. albopictus* has even been reported in regions of Spain with less than 300mm (Eritja et al., 2005). For the sake of simplicity, as the driest regions of Southeastern France (with annual rainfall close to 500mm) are already colonized by *A. albopictus* since 2010 (see Supplementary Fig. S2) we did not take rainfall into account in this study. As proposed by Waldock et al. (2013), a possible way to tackle the precipitation issue, would be to model two distinct breeding populations, that are respectively rainfall-dependent and rainfall-independent, and to use a measure of urbanisation as a proxy of the rainfall-independent breeding sites.

Regarding the management of invasive populations, the reduction of their size and the limitation of the epidemic risk of arbovirus transmission have to be envisioned in a space-time framework that can be grounded on the DISTIGRI model, which offers high spatio-temporal resolution and allows anticipation. Previous approaches involving ODE models aimed at describing *A. albopictus* abundance in South-East France based on climatic variables (Tran et al., 2013), or at designing efficient mosquito population control strategies, with sterile insect release techniques (Strugarek et al., 2019; Haramboure et al., 2020; Tran et al., 2020), but over limited geographical areas (e.g., Reunion island). The DISTRIGI model provides updated predictions by taking into account real weather data and large-scale realistic landscapes, on which efficient release strategies of sterile or Wolbachia-infected mosquitoes in a much realistic context could be designed (Dufourd and Dumont, 2012, 2013; Strugarek et al., 2018; Nadin and Toledo Marrero, 2020). Another challenging extension would be to use the DISTIGRI model to target zones and periods with high epidemiological risks, i.e., zones and periods corresponding to a high risk of outbreaks of diseases vectored by *A. albopictus*. In this case, a realistic assessment of the risk of outbreaks could be achieved by coupling the DISTIGRI model with a spatio-temporal SIR (susceptible-infected-removed) compartmental model (e.g. Pandey et al., 2013) formulated for the mosquito and human populations, which takes into account the spatial human density, the spatio-temporal variations in *A. albopictus* density and disease introduction potential (e.g., using worldwide disease patterns and airline transportation data). The high resolution of DISTIGRI and its anticipation capacity would allow to finely identify space-time windows where epidemiological surveillance and control measures should be applied.

## Declaration

### Availability of data and material

The inter-annual data on *A. albopictus* distribution are available as Supplementary Fig. S2, and from ECDC, https://ecdc.europa.eu/en/disease-vectors/surveillance-and-disease-data/mosquito-maps. The mean annual temperatures are available from Météo France: http://www.meteofrance.fr/climat-passe-et-futur/bilans-climatiques/. The temperature data at the hourly time step on a 8 × 8km^2^ grid covering France were provided by Météo France https://donneespubliques.meteofrance.fr/. Their download requires a subscription.

### Code availability

The source code for the simulation is available in the Open Science Framework repository: https://osf.io/tukgf/. The simulation requires the code to be coupled with the SAFRAN database.

## Authors’ contributions

All authors conceived the model, L.R. wrote the first draft, O.B. carried out the numerical computations, all authors discussed the results and contributed to the final manuscript.

