## Supplementary material for "Dynamics of *Aedes albopictus* invasion Insights from a spatio-temporal model": SI

#### 1 Coupled system of 2D and 1D equations

We used the 2D/1D framework developed in (Roques and Bonnefon, 2016). In this framework, the domain was divided into 2D patches (the 'matrix') surrounded by 1D edges (the 'roads'). We denote here by $a_1$ and $a_2$ the population densities on the roads and in the 2D patches ($a_2$ is simply $a$ in the main text), respectively. The dynamics in the 2D patches and the 1D edges, and the fluxes between the patches and edges are described by a system of reaction-diffusion equations. In the current framework, the intra-annual dynamics in each 2D patch $\Omega$ (see Fig. 1 in Roques and Bonnefon, 2016) can described by the following system:

$$\left\{\begin{array}{l} \partial_t w_0(t,x,y) = -\gamma_0(t,x,y)\, w_0(t,x,y) \\ \partial_t a_2(t,x,y) = D\,\Delta a_2 + a_2\, R(t,x,y,a_2) + \gamma_0(t,x,y)\, w_0(t,x,y), \\ \partial_t w_1(t,x,y) = \gamma_1(t,x,y)\, a_2(t,x,y). \end{array}\right.$$

The interactions with the roads surrounding the patch $\Omega$ are described with the boundary conditions: at each point $(x,y)$ of the boundary $\partial\Omega$ of $\Omega$,

$$D\,\nabla a_2 \cdot \nu(t,x,y) = \rho_{12}\, a_1(t,x,y) - \rho_{21}\, a_2(t,x,y),$$

with $a_1(t,x,y)$ the (linear) population density at the point $(x,y)$ of the road, $\mathbf{n} = \mathbf{n}(x,y)$ the outward unit normal to the boundary $\partial\Omega$, $\rho_{12}a_1(t,x,y)$ the flux of individuals leaving the road and entering the patch $\Omega$ at time $t$ and at the position $(x,y)$ and $\rho_{21}a(t,x,y)$ the flux of individuals leaving the patch $\Omega$ and entering a road. On the exterior boundary edges (country boarders) we assume standard reflecting boundary conditions:

$$\nabla a \cdot \mathbf{n} = 0.$$

We describe the dynamics of the population densities on the roads with 1D diffusion equations, namely, on each (linear) portion of road:

$$\partial_t a_1 = d\,\partial_{zz} a_1 + \rho_{21}\, a_2(t, x(z), y(z)) - \rho_{12} a_1(t, z) + \text{ road crossing terms.}$$

Here, the diffusion coefficient $d$ measures the mobility of the individuals on the roads. The 'road crossing terms' describe the exchanges between the two sides of the roads, are taken into account through a permeability parameter $\alpha > 0$, see (Roques and Bonnefon, 2016) for more details. Here, we assume that the permeability parameter is large enough to obtain very similar population densities on each side of each road.

The parameter values for $d$, $D$, $\rho_{12}$ and $\rho_{21}$ were estimated in (Roques and Bonnefon, 2016), based on inter-annual data and a maximum likelihood procedure. Here, we use the same parameter values, after a scaling in space: $d = 6.5 \cdot 10^6\,\mathrm{km}^2 \cdot \mathrm{yr}^{-1}$, $D = 5.4\,\mathrm{km}^2 \cdot \mathrm{yr}^{-1}$, $\rho_{12} = 4.3 \cdot 10^3\,\mathrm{yr}^{-1}$, $\rho_{21} = 2.6\,\mathrm{km} \cdot \mathrm{yr}^{-1}$.

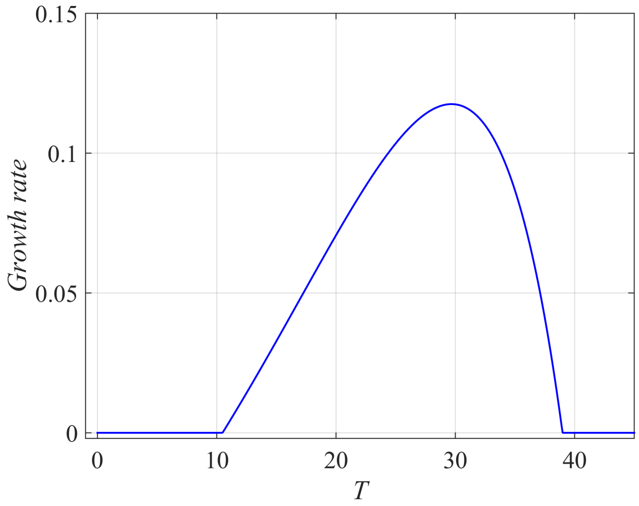


Figure S1: Growth rate $\psi(T)$ in controlled conditions with constant temperature $T$ (from Delatte et al., 2009).

### 2 Effect of the temperature on the growth rate: definition of the function $g_T$

In their study, Delatte et al. (2009) estimated the development rate (equivalently, the growth rate) of *A. albopictus* in controlled conditions, at eight constant temperatures. To obtain a value for the development rate over a full range of temperature $T \in 0-40$°C, they fitted a nonlinear model proposed by Lactin et al. (1995) to their data (fig. 1 in Delatte et al., 2009), leading to a relationship $R = \psi(T)$, with $T$ the constant laboratory temperature and $R$ the growth rate (the function $\psi$ is depicted in Fig. S1). In our case the temperature $T(t, x, y)$ is given at an hourly scale. At hourly intervals, we therefore compute $g_T$ as $g_T(T(t, x, y)) = \psi(T(t, x, y))$.

#### 3 *A. albopictus* distribution: inter-annual data

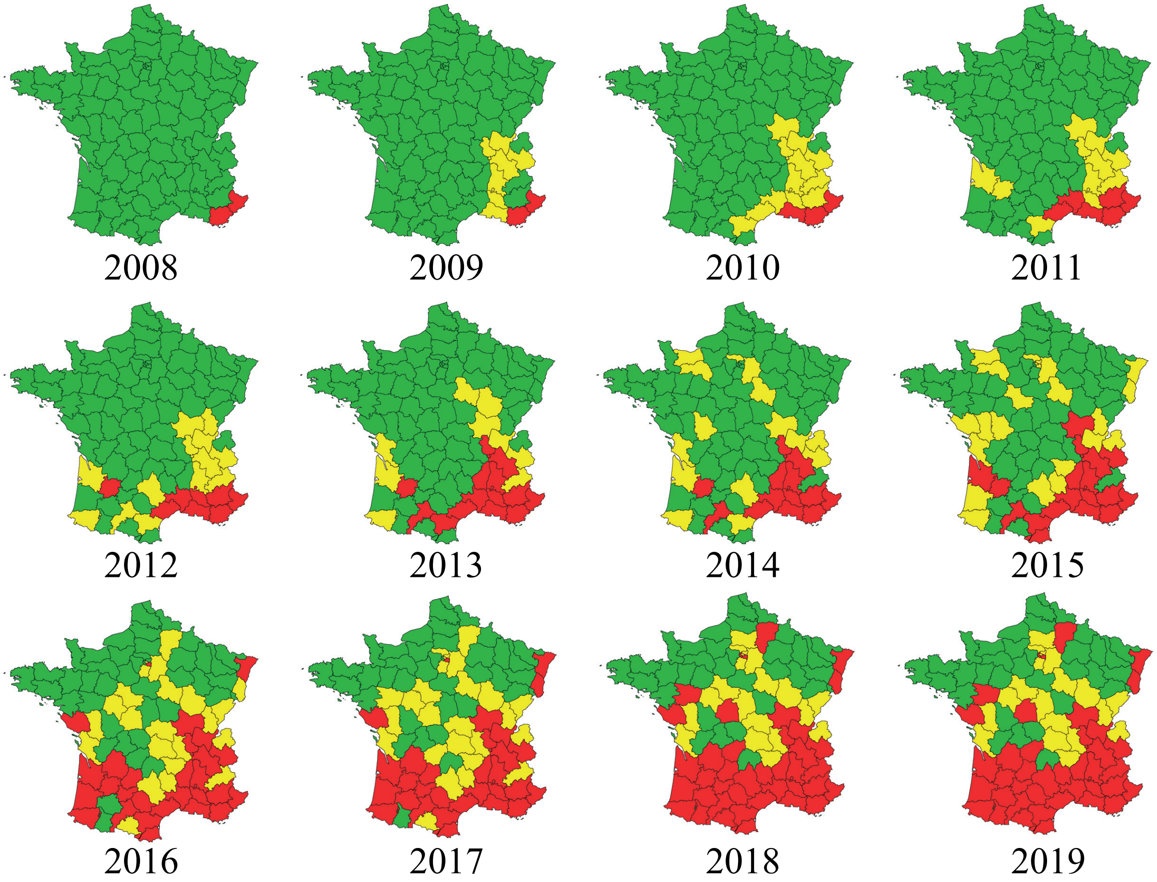


Figure S2: Distribution of *A. albopictus* in mainland France between 2008 and 2019. The mosquito was considered as absent in the green counties, introduced (few individuals detected) in yellow counties, and established in red counties. Source: French Interdepartmental Agreement for Mosquito Control.

### 4 Predicted adult density in August 2027

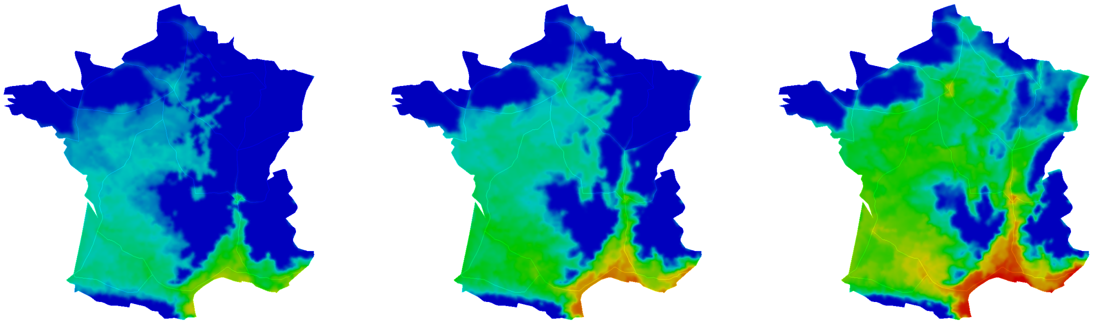

(a) 0.05-quantile 2027 (b) median 2027 (c) 0.95-quantile 2027

Figure S3: **Predicted adult density in August 2027.** At each position $(x, y)$ we show the 0.05-quantile, the median value and the 0.95-quantile of the average adult population density in August (left, central and right columns, respectively). The distribution and the quantiles are computed from 100 replicate simulations with random climatic data (selected from the 2008 − 2019 data, full year).

### 5 Predicted length of the period with significant adult occurrence

We depict in Fig. S4 the predicted duration of the period with significant adult occurrence, at each position $(x, y)$, defined here as the length of the period where $a > 0.1$. We show the 0.05-quantile (corresponding to a cold scenario), the median value (temperate scenario) and the 0.95-quantile (hot scenario), corresponding to the 100 independent climate series of the simulated climatic dataset.

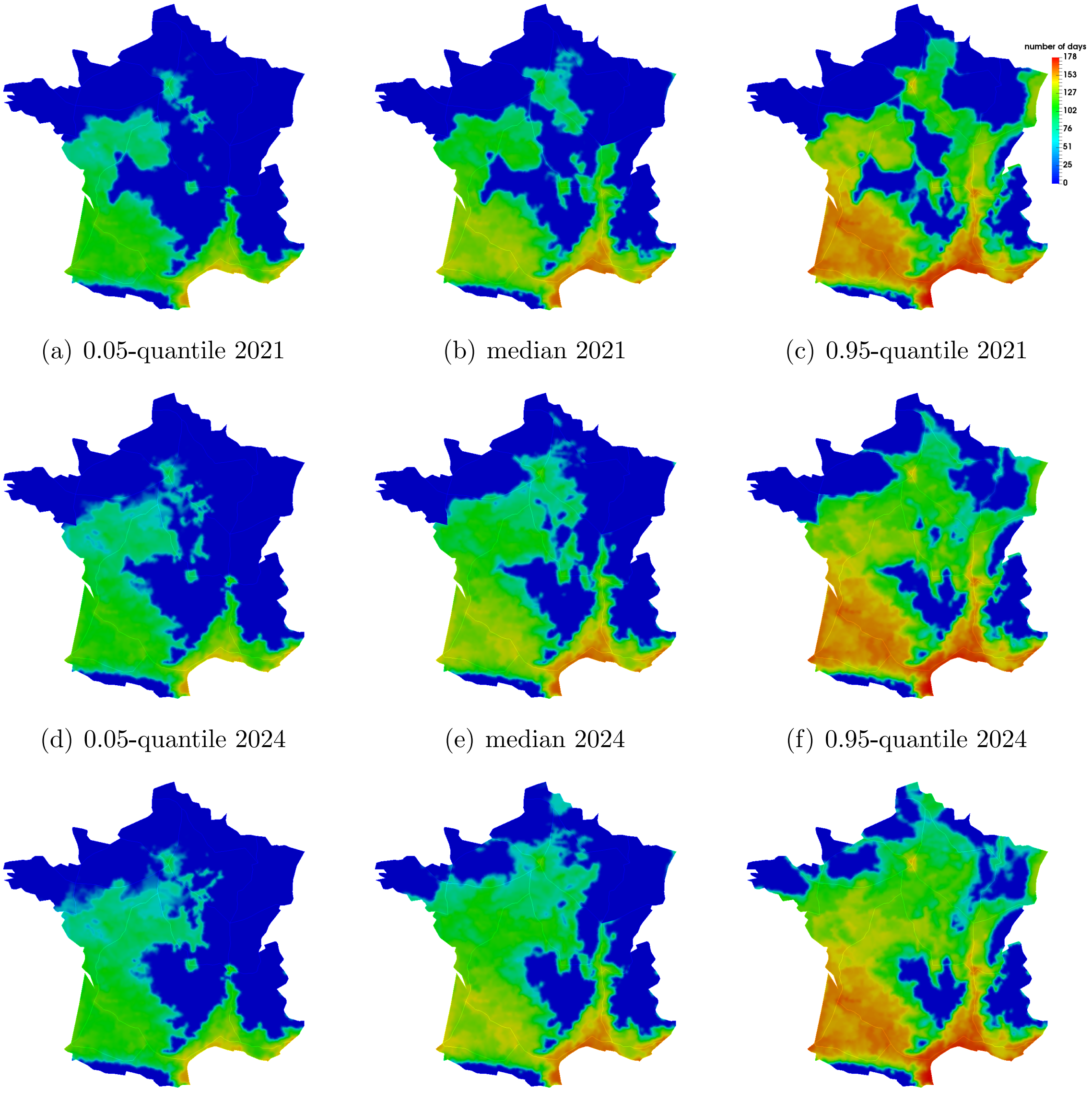


(a) 0.05-quantile 2021 (b) median 2021 (c) 0.95-quantile 2021

(d) 0.05-quantile 2024 (e) median 2024 (f) 0.95-quantile 2024

(g) 0.05-quantile 2030 (h) median 2030 (i) 0.95-quantile 2030

Figure S4: **Predicted duration of the period with significant adult occurrence over the period 2021-2030.** At each position $(x, y)$ we show the 0.05-quantile, the median value and the 0.95-quantile of the duration of the period with significant adult occurrence, defined here as the length of the period where $a > 0.1$. The distribution and the quantiles are computed from 100 replicate simulations with random climatic data (selected from the $2008 - 2019$ data, full year).
